# Antibiotic and novel compounds manipulation in vitro collagen matrix cells changes extracellular matrix non-complete cell division of fibroblast cells as new dermology technology

**DOI:** 10.1101/2021.10.07.463464

**Authors:** Waseem Ahmed, Rafia Azmat, Abdul Qayyum, Rasheed Ahmed, Sumeira Moin

## Abstract

Fibroblasts are several cells that are essential for human skin function and regulation process, the underfeed cells are a further issue of skin disorder the current study was based on isolated novel antibiotics compounds comparison of (Chloramphenicol IV) with the changes of Extracellular matrix (RC), inflammatory cells (SC) and non-complete cell division (ICD) effects on fibroblasts cell changes with the cell wall in structural and morphological changes. The new antibiotic compounds were measured and characters in (FTIR) methods with their functional group’s analysis of bioactive compounds from *Adhatoda vasica* and *Calotropis procera* plants and their effective inhibition concentrations (I C50) extract’s against tyrosinase conditions with their activity in vitro enzymatic process, both extracts have higher enzymatic inhibition assay was assessed. The fibroblast cells were compared with Chloramphenicol IV antibiotics with extracted compounds the cell wall was indiscretion and complete shape and structural changes were measured. The higher values of Diphenolase (22.5 μg/mL) was noted in *Adhatoda vasica* while an IC50 value of Monophenolase was 19.16 μg/mL, which is helpful in the treatment of fibroblast cell disorders, were higher in collagenase inhibition assay, elastase inhibition assay, hyaluronidase inhibition assay, tyrosinase inhibition assay process. It was concluded that novel antibiotics compounds from species could act an as effective role in fibroblast were used in future medicines as sources of locations and creams to control various skin diseases and skin disorder management’s processes.

## INTRODUCTION

The skin disorder is a major issue for demonologists for their prevention, the skin diseases are linked to other body disorders such as obesity, renal disorder, metabolic disorder process, the Parkinson’s and Alzheimer’s, the active compounds of many drugs founds in plants which are major secondary metabolites, the pathogenic fungi, bacteria, caused the skin disorders the fungal skin disease are most top 10 diseases are reported in globally, the enzyme inhabitation process is an important factor for change in skin structural, the old and traditional medicines are commonly used to treat the skin diseases the modern drugs used is less than 1 % (Ahmed et al. 2021). The skin is an external barrier that protects the body and has two major layers, the epidermis and dermis. The epidermis is a ubiquitous epithelium made up of keratinocytes (about 95%) and other small cell types such as melanocytes, Merkel and Langerhans cell (Ahmed et al. 2021). The dermis is the largest layer under the epidermis and plays an important role in skin formation and functioning 5.6. It consists mainly of the outer cell-matrix (ECM) produced by its many fibroblasts and includes many other cell types due to various binding structures, such as vasculature, nerves, sweat glands, and lymphatic vessels (Contra et al. 2015). Complete cell classification of all types of skin cells, as well as detailed knowledge of their functions and interactions, is important to understand the skin of homeostasis for health and disease. Fibroblasts alter the structure associated with the external environment (Momtaz et al., 2008). While sun-protected skin, fibroblasts are highly permeable, in severely damaged skin, many cells have expressed a folded / side shape. The collagen depletion is one of the hallmarks of photographic skin, the basic mechanism based on cell formation still needs to be explained in details. We investigated the morphological changes of fibroblasts caused by extracted compounds we also described a substance that protects the skin from dehydration that reduces morphological changes of skin fibroblasts through the novel antibiotic extracts on *Adhatoda vasica* belongs to the Acantaceae family, commonly known as Malabar nut plants, and contains activators of phytochemicals. *Colotropis procera* is a flowering plant belonging to the family Assiliapedesi (Lima) (Ahmed et al. 2018). It is widely distributed in Asia and the subcontinent as well as other parts of West Africa and tropical Irvine. Antioxidant bioactive compounds are widely distributed in plants reported to have many biological effects, including antioxidant, free radical scavenging, anti-inflammatory, anti-carcinogenic Carvalho et al. 2005. (Ahmed et al. 2018) reported skin problems are a major problem in the world. Tyrosine activity is a problem of the skin of the eye and many other disorders (Contra et al. 2015).

The research article discloses the isolation often essential antibiotic compounds from *Adhatoda vasica* and *Calotropis procera* using UF-HPLC-DAD and Fourier transform infrared spectroscopy (FTIR) analysis of new compounds and the morphological changes of fibroblast under the changes of cells were completely measured in election transmission electron scope techniques with linked to with the potential activity of diphenolase and monophenolase for checking the activities of new compounds and inhibition assay study (collagenase inhibition, elastase inhibition assay in plant exacts. The complete comparison of Extracellular matrix (RC), inflammatory cells (SC) and non-complete cell division (ICD) effects on fibroblasts cell changes under antibiotic and novel compounds changes. The liquid exacts were apply on skin cells of fibroblast and their changes were measured through morphological characterization of fibroblasts skin cells, the new compounds were used as future green resolution, as a drugs preparation and developmental process in various pharmacy and pharmaceutical industries uses as drug manufacture of skin diseases.

## 2. EXPERIMENT

### Collection of herbal plants and experimental site

Fresh mature leaves were collected from Khanpur Valley, Haripur, Pakistan and transferred to the Horticulture Laboratory University of Haripur for analysis at the Department of Chemistry, and the University of Karachi, which assisted in the chemical isolation of the bioactive compound. The coordinates of the collection points at Haripur are 33.9946 N, 72.9106 E.

### UF-HPLC-producer accuracy methods of the compounds isolation and measurement process

The testing process has been used based on previous research and consists of three stages namely incubation, washing and isolation. The tested SMD solution was briefly set to “100” “L” 200 L HSA (600 buffer) and the buffer solution “200” L was set at 37 ° C for 20 min. Meanwhile, the HSA solution was boiled for 15 seconds. With a fixed weight molecule of 30 kDa (Millipore Amicon Ultra-0.5 ml, item, UFC 503096) and 14,000. Separate unspecified materials from HSA-ligand structures for 15 minutes at room temperature. The residue was washed with 200 cm centrifugation to remove the unbound material. At HSA they were released to the assembly with an elucidation of 400 mL 50% methanol (pH = 3) for 20 min and then centrifuged at 14,000 × g for 15 min at room temperature, which is a dual process. Separate the fillets, and then the joint edition. And 1000 “μL” were added by 50% methanol and then directly analyzed (Da et al. 2016).

### Fourier transform infrared spectroscopy (FTIR) analysis for the compounds analysis process as fibroblast disorder management process

Fourier transform infrared spectroscopy FTIR (Model / Mac25 Germany) with attached data software and operating procedures. A small amount of all the flower buds and seeds of both plants were prepared from Kell pellets for FTIR analysis and a small pressure film was done. All samples were analyzed three times with KBR blank. Spectral data were compared to identify functional groups present in the samples (Ahmed et al. 2019).

### Tyrosinase inhibition property of active compounds for fibroblast diseases

Tyrosinase inhibits the property of active compounds for skin diseases was determined by the mechanism of action of tyrosinase inhibition (Khan et al., 2012). Two herbaceous plant samples were calculated with the help of formula percentage, thereby; % Tyrosinase inhibition = [(one control - one pattern) / one control] x 100.

### Enzymatic and inhibition procedures for fibroblasts skin cells treatments with enzymatic assays

### Collagen inhibition assay for fibroblast

Collagenase resistance testing has been described as a method based on the hydrolysis of N- [3-[3-(2-furyl)acryloyl]-Leu-Gly-Pro-Ala (FalgPA) by minimally altered collagenase. Collagenase 511 mM NaCl and 21 mM CaCl2 (pH 7.5) from Clostridium histolyticum (COL) were dissolved at 0.9 units / mL substrate in the buffer, at FALGPA, and 60 mM trisin buffer was used. Trisin buffer up to 2 mm. The sample extract was kept with the enzyme in a bath for 17 min before adding the surface to the reaction. Tricin buffer, 0.9 mM FALGPA, 0.1 units of ChC and a final reaction mixture (total 150 μl volume) containing 25 g for experimental extraction. While the control work was carried out by water and methanol, oleanolic acid was used as good control, after injecting the substrate for 20 min, the collagenase activity was measured at 340 nm.

### Elastase inhibition assay methods for fibroblast

Elastase inhibition has been attributed to the use of N-suck- (ala) 3-nitronilide (SANA) as a substrate to monitor the release of the enzyme p-nitroaniline Gulluce et al., 2006). The resistivity was determined by the strength of the pigment extracted from the SANA crack by the elastase reaction, adjusted with the SANA 1 mm 0.2 mM Tris buffer (pH 8.0). The solution was added to a vortex at 26 C for 13 min and then elastase was added to a 0.02 / ml unit. The solution was vortexed in a hot water bath at 26 ° C for 12 min and the absorption is measured at 411nm. Methane and water were regulated when positive oleanic acid was used.

### Hyaluronidase activity for fibroblast cells of skin

The hyaluronidase inhibitory action of the test was modified with some modifications to the previously described method Ahmed et al. 2019. The bovine test was carried out by adding 2.0 units per 100 units of sodium phosphate buffer 22 mm (pH 7.0) serum albumin with bovine 78 mm (BAS) 0.02% at 37 C for 11 minutes by adding seed comb and hyaluronic acid. Sodium salt at 38 ° C 200 l (0.03% in 300 mM, pH 5.35) Start incubation Albumin in sodium acetate Serum Albinic hyaluronic acid (not ground) 1 ml Albumin acid 0.2% dissolved in 41 minutes with the addition of acetic acid at 25 mm and 80 mm, Measured at 600 nm. The oleanolic acid was well regulated by the sample used in this study. Percentage resistance to collagen, elastase and hyaluronidase tests is calculated by: Enzyme inhibition function (%) = (1– b / a) x 100, where A is the enzyme activity outside the sample and B is the function in the sample.

### NADH System Superoxide-Radical Scavenging Assay

Superoxide release was measured according to the method of Lae et al with some changes. The mixture contained a 0.6 ml sample of 105 mM NADH, 0.5 ml of 66 mM NBT in a 0.1 M phosphate buffer (pH 7.4) and 0.1 ml at various concentrations. Water and methanol were used as regulators and BHT as good regulators. After 10 minutes, the reaction mixture reached a consistent colour, with an absorption rate of 560 nm.

Power of DPPH, ABTS · +, *** - calculated according to NADH-based formula:

Extinction (%) = [(O.Dcontrol - O.Dsample) / O.Dcontrol] x 100, Where O D is the visible quantity, where there are samples with or without samples.

Determining the effects of peroxinitrite mutationsDetermining effects on peroxinitrite changes

### Determination of Peroxynitrite changes

Peroxynitrite was compounded by the method of Beckman et al. [17]. The acidic (0.6-M HCl) solution of 6-mL H2O2 was mixed with 5-mL 0.6 M KNO2 in a 1.3-M NaOH ice bath added to the reaction component. Excess H2O2 was removed by granular MnO2 treatment with 1.3-M NaOH and the reaction mixture was left overnight at −20 ° C.

### Lactate dehydrogenase (LD or LDH)

The cytotoxicity of both herbals plants samples was examined using a Kit of LDH. The extract of (2 ml) both herbal plants samples were taken where the release of lactate dehydrogenase (LDH, cytoplasmic enzyme) treated with the cells was determined according to the calorimetric method as reported by (Muzitano et al., 2006). The different concentrations were used (1, 0.4 and 8 g/ml) and setting the sample for 24 h. The lysates were obtained by application of 1 % X-100 Trition solution with a similar ratio of DMSO solution. The process was further diffused and compared with control samples.

### Scanning electron and microscopy studies of fibroblast of skin under relation of antibiotic and novel compounds

Skin cells enlarged by collagen gels are purified with PBS and prepared in 4% glutaraldehyde (Wako, Osaka, Japan) in PBS for 24 hours scanning electron microscopy (SEM). After preparation, these cells are first immersed in 1% osmium tetroxide (Wako) for 1 hour. Following the dehydration of ethanol, the cells were at a critical point dried with liquid CO2, injected into SEM stubs with carbon tape and sprayed with gold. Samples were viewed and photographed with a scanning electron microscope (JSM-6701F, JEOL, and Tokyo, Japan). By measuring the area of the horizontal cell, the HDFs produced in the vessels are encased in crystal violet and photographed with a simple microscope (BX51, Olympus, and Tokyo, Japan). The location of the individual cell cells was mass-tested using the NIH Image software.

#### Morphological characterization of fibroblast cells changes extracellular matrix (RC), inflammatory cells (SC) and non-complete cell division (ICD)

The cells of fibroblast with their morphology the changes of adhesion, spreading were examined the cells and their morphology change adhesion, diffusion was tested for 30 min, 4 and 24 h, as described by de Oliveira and Nanci (7). Briefly, cells sown in glass coverslips were prepared using 4% paraformaldehyde at 0.1 M sodium phosphate buffer (PB), pH 7.2, 10 min. After that, they were filled with 0.5% Triton X-100 in PB for 10 minutes. Alexa fluor 488 (green fluorescence) - combined phalloidin (1: 200) (Molecular Probes, Invitrogen, OR, USA), and 300 nM 4’, 6-diamidino -2-phenylindole, dihydrochloride (DAPI, Molecular Probes) was used to locate the actin cytoskeleton and cell nucleus. After incorporation and anti-corrosion kit (Vectashield; Vector Laboratories, Burlingame, CA, USA), the samples were tested under epifluorescence.

## 3. STATISTICAL ANALYSIS

Details are presented here as ± SEM. All data were analyzed for differences between the tested methods using the Duncan test. The difference was considered significant at p <0.05. All analyzes were performed using Graph Pad software, version 12.01 (USA). Chemicals are guaranteed to be developed and validated and measure the quality of chemicals by digital libraries. HPLC-DAD data was measured with the help of Chrom Gate v 3.31 Knauer software used (Ahmed et al. 2021).

## 4. RESULTS

### The Novel antibiotic compounds from herbal plants as management of fibrobalsting skin cells

The new active compounds of Viscic acid, Pasakbumin C, 8’Z-enyl congener, 5-(8’Z,11’Z-heptadecadienyl)-1,3 benzenediol, 9’-(o-methyl) protocetraric acid and Calophynic acid isolated from *Adhatoda vasica* and *Calotropis procera* is reported in Table 1. The higher contents of 1-(8’Z, 11’Z-heptadecadienyl) (15 mg/100g) were found in *Adhatoda vasica* leaves whereas lower contents were measured in *Calotropis procera*. However, the contents of <u>Viscic acid</u> were 13 mg/100g and (11 mg/100g) in leaves of *Adhatoda vasica* and *Calotropis procera*, same the as all other compounds were higher in *Adhatoda vasica leaves* as compare to *Calotropis procera*. The 8’Z,11’Z-heptadecadienyl was higher in the seed of both plants while the other contents were higher in *Calotropis procera*. The chorogram of all peaks of new compounds were shown in figure 3.

**Table 1.** UF-HPLC isolation of new compounds in herbal plants as skin fibroblast disorder managements

| Novel antibiotic compounds (mg/100g) | Herbal plants |  |  |  |
| --- | --- | --- | --- | --- |
|  | *Adhatoda vasica* leaves | *Calotropis procera* leaves | *Adhatoda vasica* seed | *Calotropis procera* seed |
| Viscic acid | 13 ± 4.51 | 11 ± 351 | 10 ± 5.51 | 11 ± 5.51 |
| Pasakbumin C | 11 ± 3.51 | 12 ± 451 | 9 ± 5.51 | 8 ± 5.51 |
| 8'Z-enyl congener | 14 ± 2.51 | 13 ± 3.51 | 11 ± 5.51 | 10 ± 5.51 |
| 5-(8'Z,11'Z-heptadecadienyl)-1,3-benzenediol | 15 ± 2.51 | 14 ± 351 | 12 ± 5.51 | 12 ± 5.51 |
| 9'-(o-methyl) protocetraric acid | 12 ± 3.51 | 11 ± 3. 51 | 11 ± 5.51 | 12 ± 5.51 |
| Calophynic acid | 10 ± 1.51 | 9 ± 4.51 | 8 ± 5.51 | 8 ± 5.51 |

± SD values with the means values for three triplicate was observed

**Fig1.**
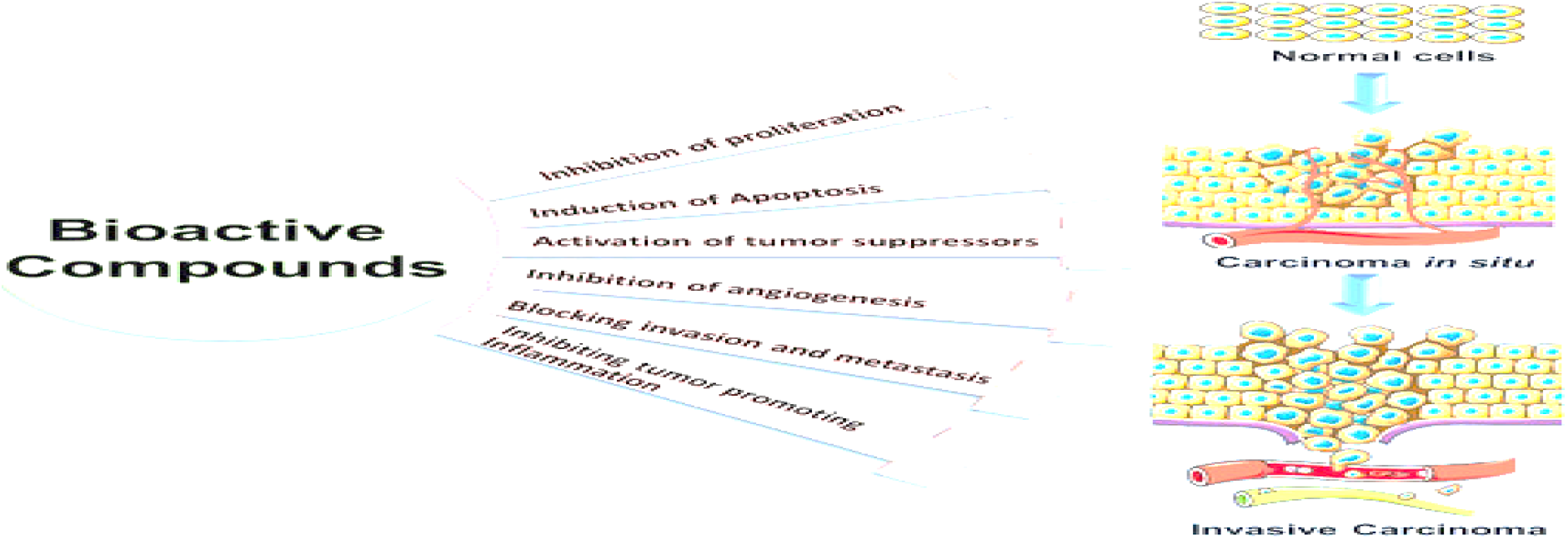
The bioactive compounds machism to control skin disasesse and its magements porcees

**Fig 2.**
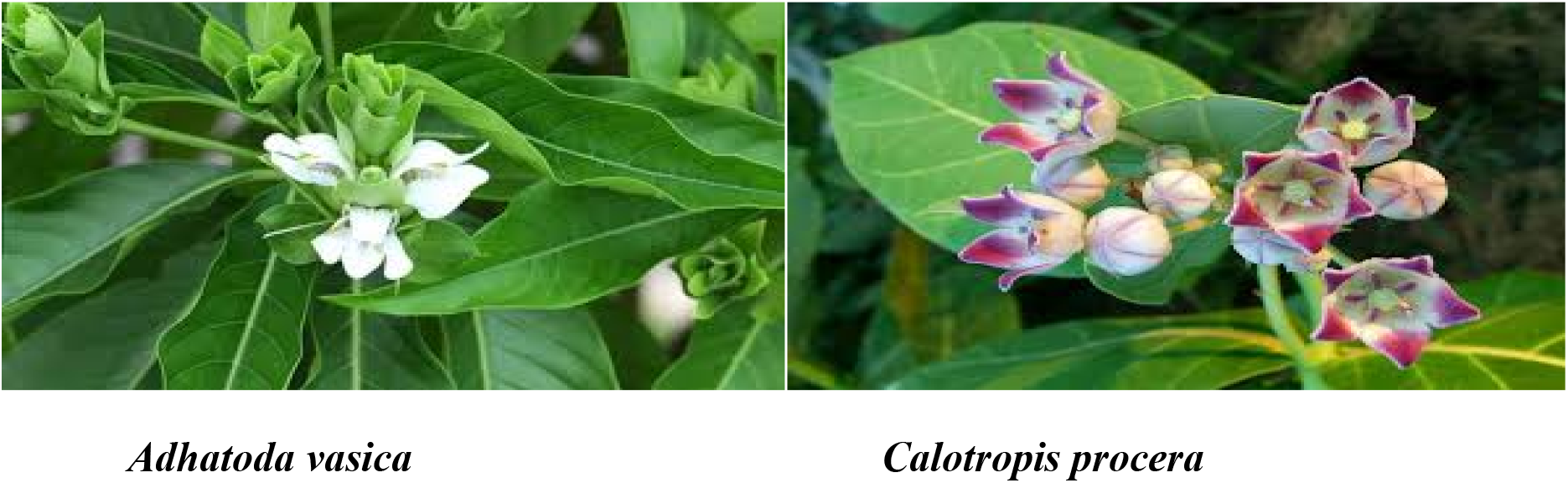
The medical plants and their parts for new medicnes

**Fig 3.**
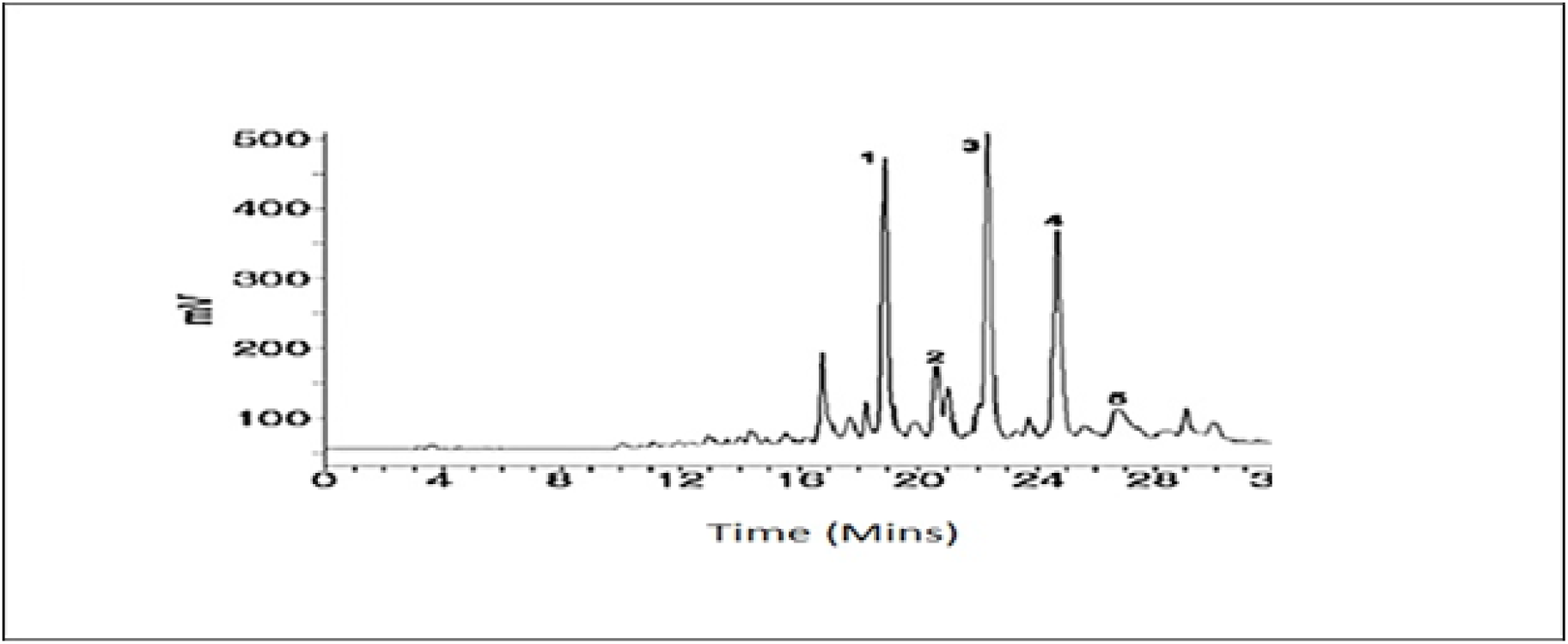
Chorogram of New compounds (1-5) Viscic acid, Pasakbumin C, 8’Z-enyl congener, 5-(8’Z,11’Z-heptadecadienyl)-1,3-benzenediol, 9’-(o-methyl) protocetraric acid and Calophynic acid

### Structural and molecular response of new bioactive compounds Fourier transform infrared spectroscopy (FTIR)

The important parameters of compounds of molecular formula, molecular weight % and peaks structure through Fourier transform infrared spectroscopy (FTIR) was performed on two herbaceous plants to identify the active group, the collected samples were analyzed by IR based on them and the results showed the presence of groups described in Table 2 and all individual peaks of all compounds were shown. All five compounds differ in molecular formula and weight as shown in table 2. The composition formula differs in terms of the highest values of the five compounds.

**Table 2.**
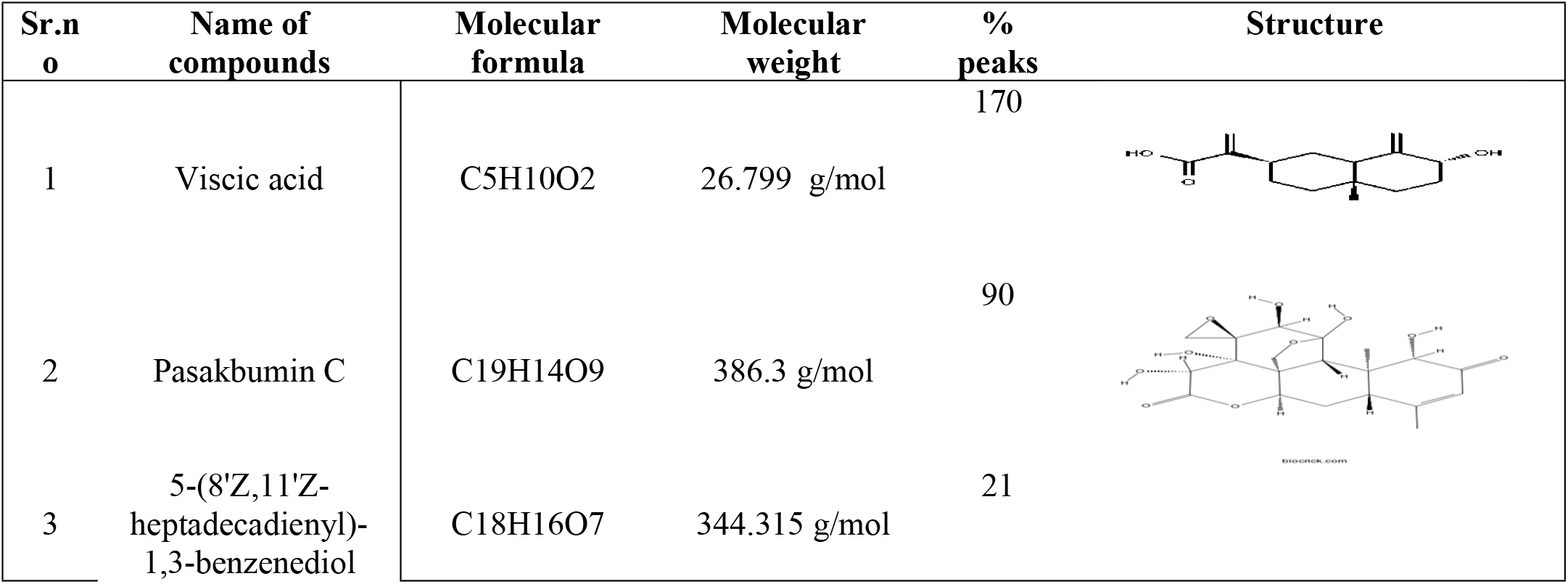
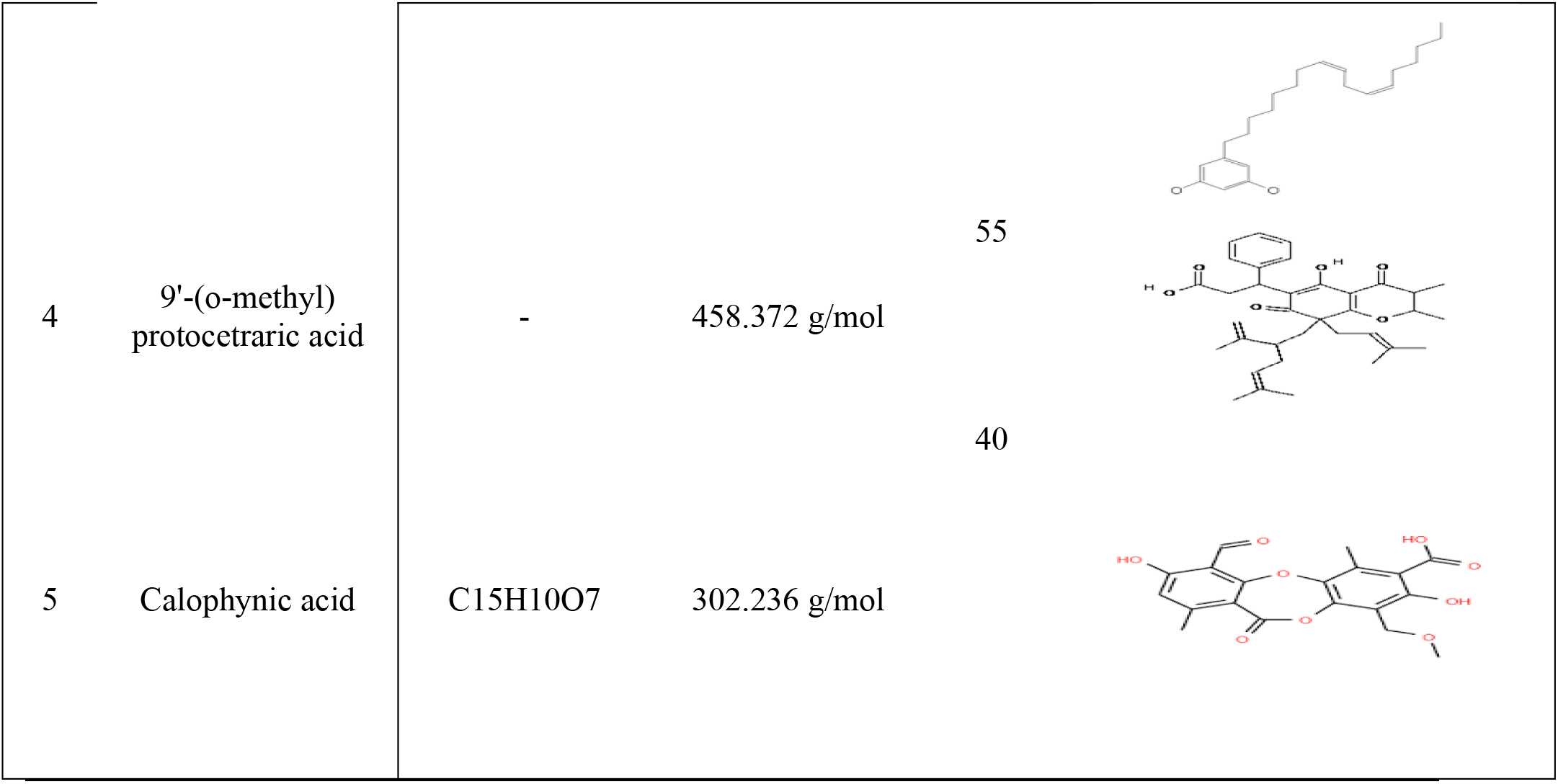
UF-FIR analysis of new compounds in herbal and comparative analysis

| Sr.no | Name of compounds | Molecular formula | Molecular weight | % peaks | Structure |
| --- | --- | --- | --- | --- | --- |
| 1 | Viscic acid | C<sub>5</sub>H<sub>10</sub>O<sub>2</sub> | 26.799 g/mol | 170 |  |
| 2 | Pasakbumin C | C<sub>19</sub>H<sub>14</sub>O<sub>9</sub> | 386.3 g/mol | 90 |  |
| 3 | 5-(8'Z,11'Z-heptadecadienyl)-1,3-benzenediol | C<sub>18</sub>H<sub>16</sub>O<sub>7</sub> | 344.315 g/mol | 21 |  |
| 4 | 9'-(o-methyl) protocetraric acid | - | 458.372 g/mol | 55 |  |
| 5 | Calophynic acid | C<sub>15</sub>H<sub>10</sub>O<sub>7</sub> | 302.236 g/mol | 40 |  |

### Functional groups analysis of new bioactive compounds by Fourier transform infrared spectroscopy (FTIR)

An important functional analysis of the various components of the two herbal plants was studied and their characteristics are presented in Table 3. Alcohol extension with Intermolecular OH Cells and extension alcohol-free OH and 3982.14 1 3.51. 322.14 of *Adhatoda vasica*. 2.51 Leaves, low values of the same parameters are shown in the seed of Calotropis procera. Intermolecular Bonded Alcohol OH extension and intramolecular bond alcohol OH reduction 3285.14 .1 5.51 3185.14 1 5.51. 51 Adatoda vasika leaves, low values of the same parameters are shown in the seeds of Colotropis procera. Simple CC alkanes and fragrant C-being compounds in the leaves of *Adhatoda vasica*. exceed 3285.14.1 5.51 3385.14 1 5.51, although the flowers and shoots indicate the values of the second and third stages. C = N Reducing Emin / Oxim or C = O Extending Combined Ketones or Alkanes 3915.14 .1 5.51, OH Bending Penol 3915.14 1 5.51 CN Stretch Amine 3935.14 ± 5.51 after lower values of flow branches and three functions

**Table 3.** Functional groups analysis of bioactive compounds by Fourier transform infrared spectroscopy (FTIR)

| Functional groups | Leaves extracts<br>*Adhatoda vasica* | Seeds extracts<br>*Calotropis procera* |
| --- | --- | --- |
| Free alcohol O-H Stretching $\text{cm}^{-1}$ | $3982.14 \pm 3.51$ | $3085.14 \pm 4.51$ |
| Intermolecular bonded alcohol O-H Stretching $\text{cm}^{-1}$ | $3272.14 \pm 2.51$ | $2985.14 \pm 3.51$ |
| Intramolecular bonded alcohol O-H Stretching $\text{cm}^{-1}$ | $3181.14 \pm 5.51$ | $2685.14 \pm 3.51$ |
| C-H stretching alkane $\text{cm}^{-1}$ | $3282.14 \pm 3.51$ | $2285.14 \pm 4.51$ |
| C-H bending aromatic compounds overtone $\text{cm}^{-1}$ | $3381.14 \pm 4.51$ | $2785.14 \pm 4.51$ |
| C = N stretching imine/oxime or C = O stretching conjugated ketone or alkenes $\text{cm}^{-1}$ | $3916.14 \pm 3.51$ | $3015.14 \pm 4.51$ |
| O-H bending phenol $\text{cm}^{-1}$ | $3911.14 \pm 2.51$ | $2815.14 \pm 3.51$ |
| C-N stretching amine $\text{cm}^{-1}$ | $3931.14 \pm 3.51$ | $2735.14 \pm 2.51$ |

### Tyrosinase inhibition activity and process

The potential activity of tyrosinase inhibition activity of these species was also monitored and reported in Table 4 where a significant difference was found in tyrosinase inhibition (IC50) at (*p*<0.05). The leaves of *Adhatoda vasica* showed the range of diphenolase and monophenolase (22.5±0.07 and 19.16±0.07 μg/mL) while *Calotropis procera* showed the diphenolase and monophenolase (15.66±0.11 and 13.66±0.11 μg/mL) respectively, in the seed of diphenolase and monophenolase (20.5±0.03 and 12.16±0.07 μg/mL) from *Adhatoda vasica* and (19.66±0.11 and 16.66±0.12 μg/mL) from *Calotropis procera* which indicated that both herbal plants have the potential activity of Tyrosinase inhibition for skin diseases and they’re infectious.

**Table 4.** Effective inhibition concentrations (I C50) extract’s against tyrosinase conditions for control of skin diseases

| Herbal Extracts | IC50<br>Monophenolase<br>(µg/mL) leaves | IC 50 Diphenolase<br>(µg/mL) leaves | IC50<br>Monophenolase<br>(µg/mL) seeds | IC 50<br>Diphenolase<br>(µg/mL) seeds |
| --- | --- | --- | --- | --- |
| *Adhatoda vasica* | 22.16±0.07a | 15.5±0.11<br>a | 20.17±0.03a | 19.5±0.11<br>a |
| *Calotropis procera* | 19.66±0.12b | 13.66±0.12 b | 12.66±0.11b | 16.66±0.12 b |
| Significant level | <0.00001 | <0.00001 | <0.00001 | <0.00001 |

Mean values in the same column followed by different letters represent statistical different ( $P<0.0001$ ) using the Duncan's multiple range test.

### Anti-collagenase and anti-elastase activities of herbal extracts as skin disorder managements

The collagenase and elastase inhibition effects of both herbal plants extracts at the final concentration of 2 mg/ml were determined and elucidated as shown in Table 5. The higher activity of collagenase 24.16±0.01 was noted in leave extracts of *Adhatoda vasica* and *Calotropis procera* also employed the 20.66±0.11 was observed in leave extracts, the anti-elastase activity was higher in *Adhatoda vasica* leave extracts, same the good activity of anti-elastase was observed in 15.61±0.11 in leaves of *Calotropis procera*.

**Table 5.** Effective inhibition concentrations of various enzymatic process for skin diseases control process

| Herbal Extracts | Collagenase<br>inhibition assay<br>% | Elastase inhibition<br>assay<br>% | Hyaluronidase<br>inhibition assay<br>% | Tyrosinase<br>inhibition assay<br>% |
| --- | --- | --- | --- | --- |
| *Adhatoda vasica* | 24.16±0.01a | 18.5±0.11a | 21.17±0.01 | 20.5±0.12<br>a |
| *Calotropis procera* | 20.66±0.11b | 15.61±0.11 b | 14.66±0.11b | 16.66±0.12 b |
| Significant level | <0.00001 | <0.00001 | <0.00001 | <0.00001 |

Mean values in the same column followed by different letters represent statistical different ( $P<0.0001$ ) using the Duncan's multiple range test.

### Anti-hyaluronidase activity in herbal extracts

The inhibitory effects of both herbal plants were evaluated in Table 5. The higher activity of Anti-hyaluronidase 20.17±0.03 was noted in leave extracts of *Adhatoda vasica* and *Calotropis procera* also employed the 12.66±0.11b was observed in leave extracts, the activity of inhibitory and velocity was higher in *Calotropis procera* as compare to the *Adhatoda vasica* shown in figure 6.

### PMS-NADH system superoxide-radical and peroxynitrite activity in leave extracts

The PMS-NADH system of superoxide-radical action was measured in both herbal and shown in table 6. The process of the PMS-NADH system of superoxide-radical action was measured in *Calotropis procera* 17.66±0.12b as compare to *Adhatoda vasica* leave extracts the activity of seed was performed higher in both extracted samples.

**Table 6.** Effects of PMS-NADH System Superoxide-Radical and Peroxynitrite leaves

| Herbal Extracts | PMS-NADH<br>system superoxide-radical leaves | PMS-NADH<br>system superoxide-radical seeds | Peroxynitrite<br>activity leave | Peroxynitrite<br>Seeds activity |
| --- | --- | --- | --- | --- |
| *Adhatoda vasica* | 21.16±0.07a | 12.5±0.11<br>a | 10.17±0.03a | 8.5±0.11 a |
| *Calotropis procera* | 17.66±0.12b | 11.66±0.12 b | 11.66±0.11b | 9.66±0.12 b |
| Significant level | <0.00001 | <0.00001 | <0.00001 | <0.00001 |

Mean values in the same column followed by different letters represent statistical different ( $P<0.0001$ ) using the Duncan's multiple range test.

### Responses of extracts under the fibroblast cells on Lactate Dehydrogenase (LD) activity

The cytotoxicity test was analyzed only in methanol extracts of both plants by using three concentrations (1, 0.4 and 0. 8). It was observed that a higher cytotoxic value for 0.8 % concentration was 87 ug/ml cytotoxicity in menthol solution. It was interesting to note that higher cytotoxicity test only measures with concentration 0.8 % as compared to 0.4 and 1 percentage solution, from isolated compounds of *C. procera* leaves extracts the maximum range of cytotoxicity level was 9087 ug/ml from 1% concentration of basic methanol extracts solutions. The however lower level of cytotoxicity was found from 0.4 and 0. 8 % solutions. The cytotoxicity values were shown in figures 4, 5. According to standardization of concentration for cytotoxicity test, it was 1% of both herbal plants.

**Fig 4.**
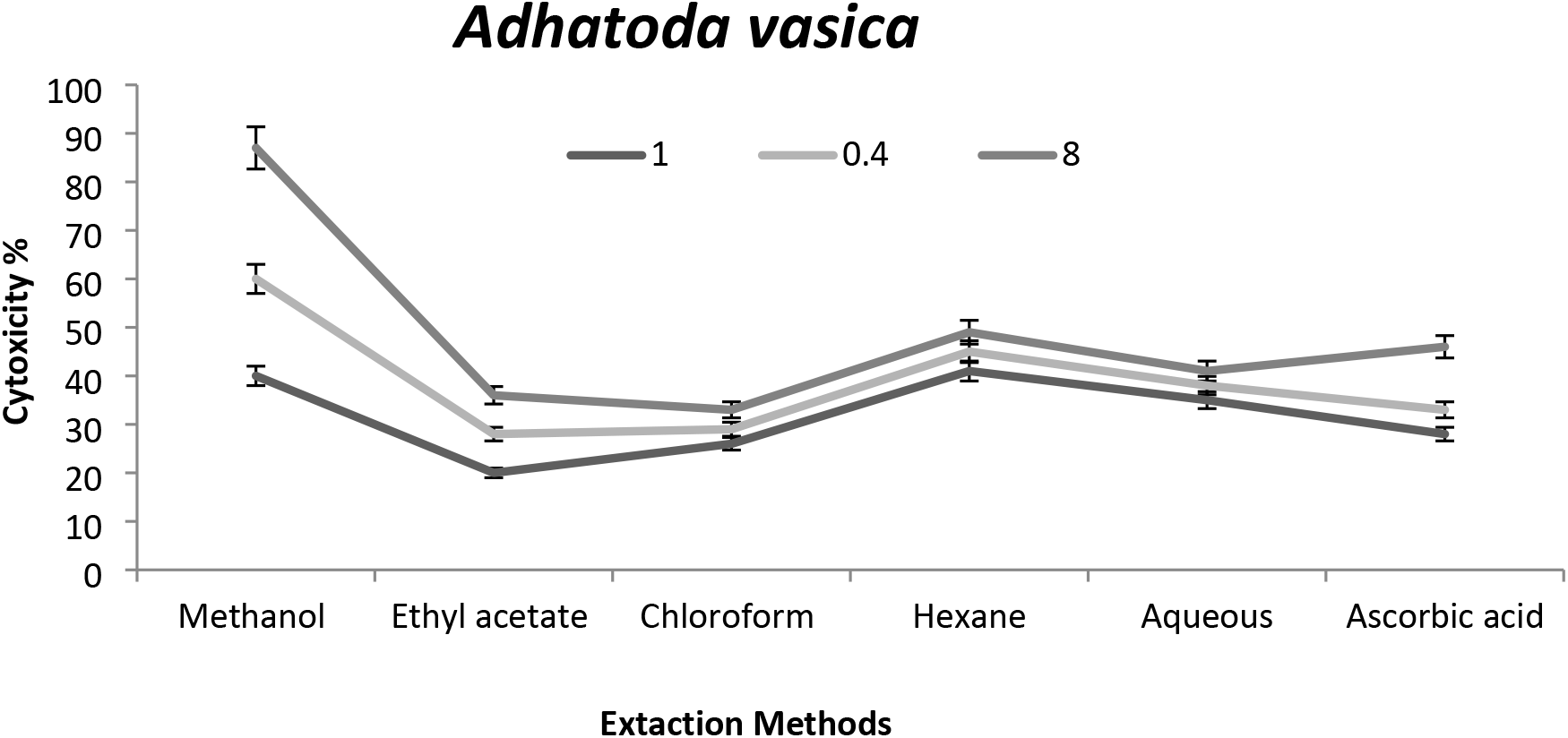
Cytotoxicity of extracts under the diverse solvent process for Adhatoda vasica

**Fig 5.**
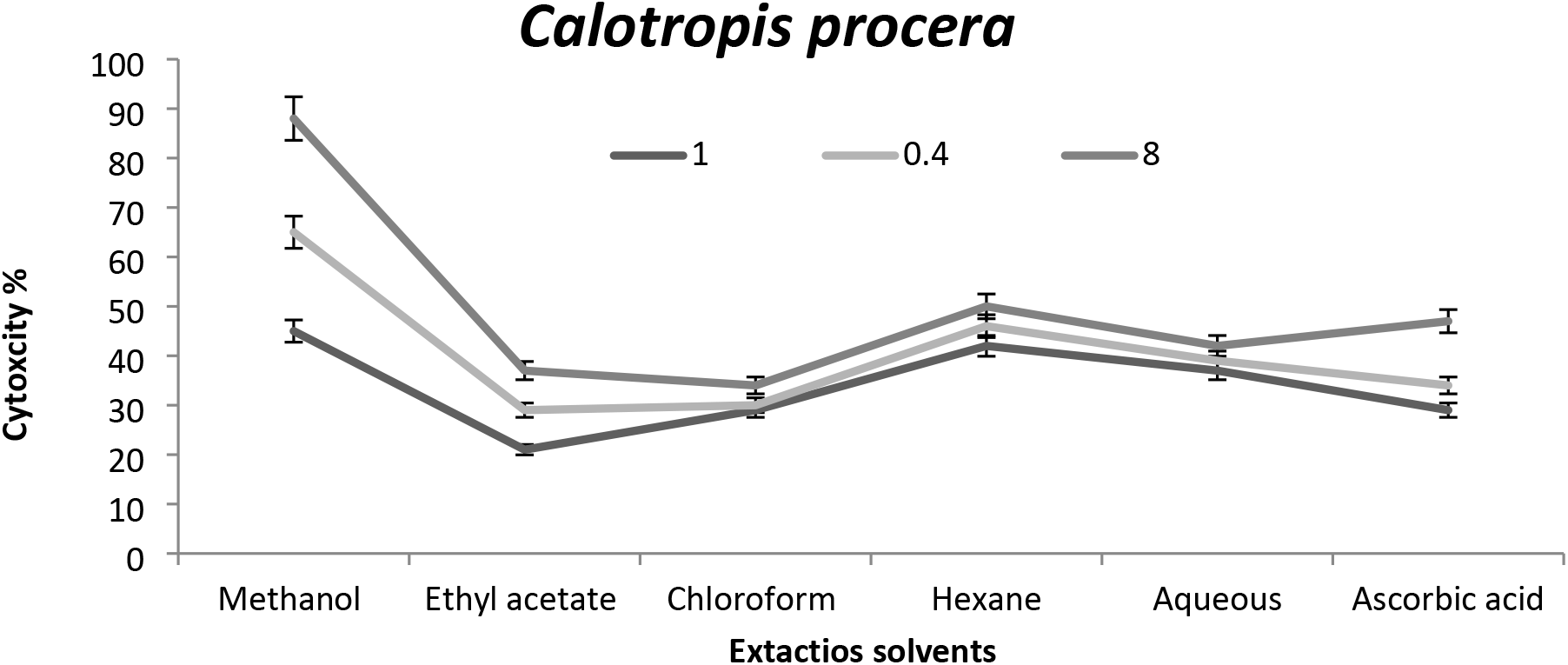
Cytotoxicity of extracts under the diverse solvent process Calotropis procera

**Fig 6.**
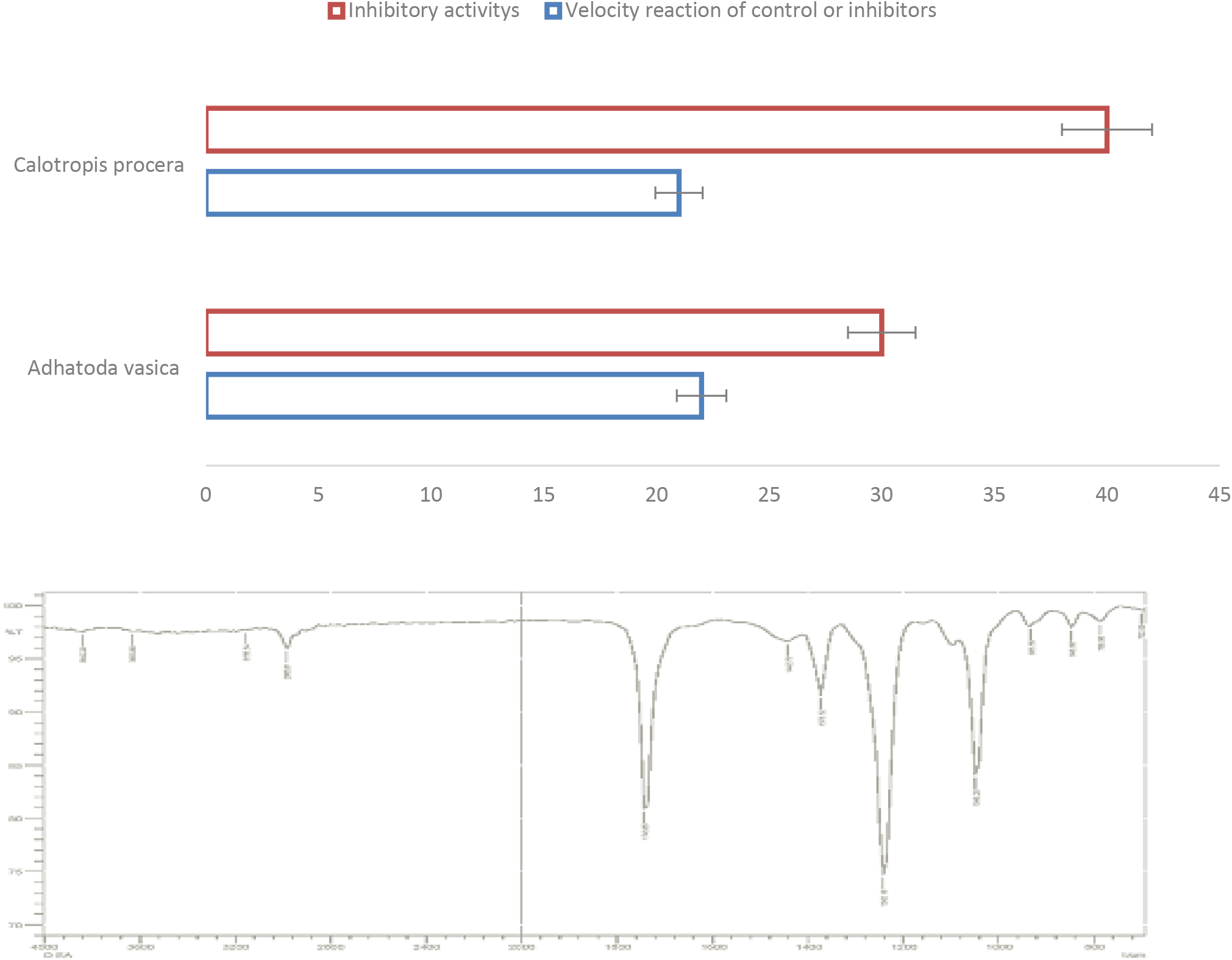
Functional groups analysis of bioactive compounds by Fourier transform infrared spectroscopy (FTIR)

### Extracellular matrix (RC), inflammatory cells (SC) and non-complete cell division (ICD) effects on fibroblasts cell changes

To evaluate the effects of liquid extracts of both herbal plants on fibroblast cells were study shown in figure 7 f any, of the oxidant / antioxidant imbalances evidenced by skin fibroblasts, cellular status was assessed by looking at two common genetic factors: natural distribution and collagen distribution of cells investigating the transfer of vital electron microscopes found in skin cells. Significant differences in endoplasmic reticulum cisternae were found in skin fibroblasts, as well as evidence of multilamellar cytoplasmic cells found in cells. This figure was shown by changes in the figure a1- d1 after measurement of extraction activity of fibroblast cells with antibiotic was a1-d1 and antibiotic compounds were a2-d2 i, the formation of perinuclear granules were found. Similarly, fibroblast cells were treated with good control (Chloramphenicol IV,) at 0.020 mg/ml and showed changes in a bacterial structure similar to those seen in cells treated with extracellular matrix (RC), inflammatory cells (SC) and non-complete cell division (ICD).) (Fig.). Both plant extracts have been shown to have significant structural changes in the cell wall and enter the structure rather than with clear exposure to cell death and fibroblast control

**Fig 7.**
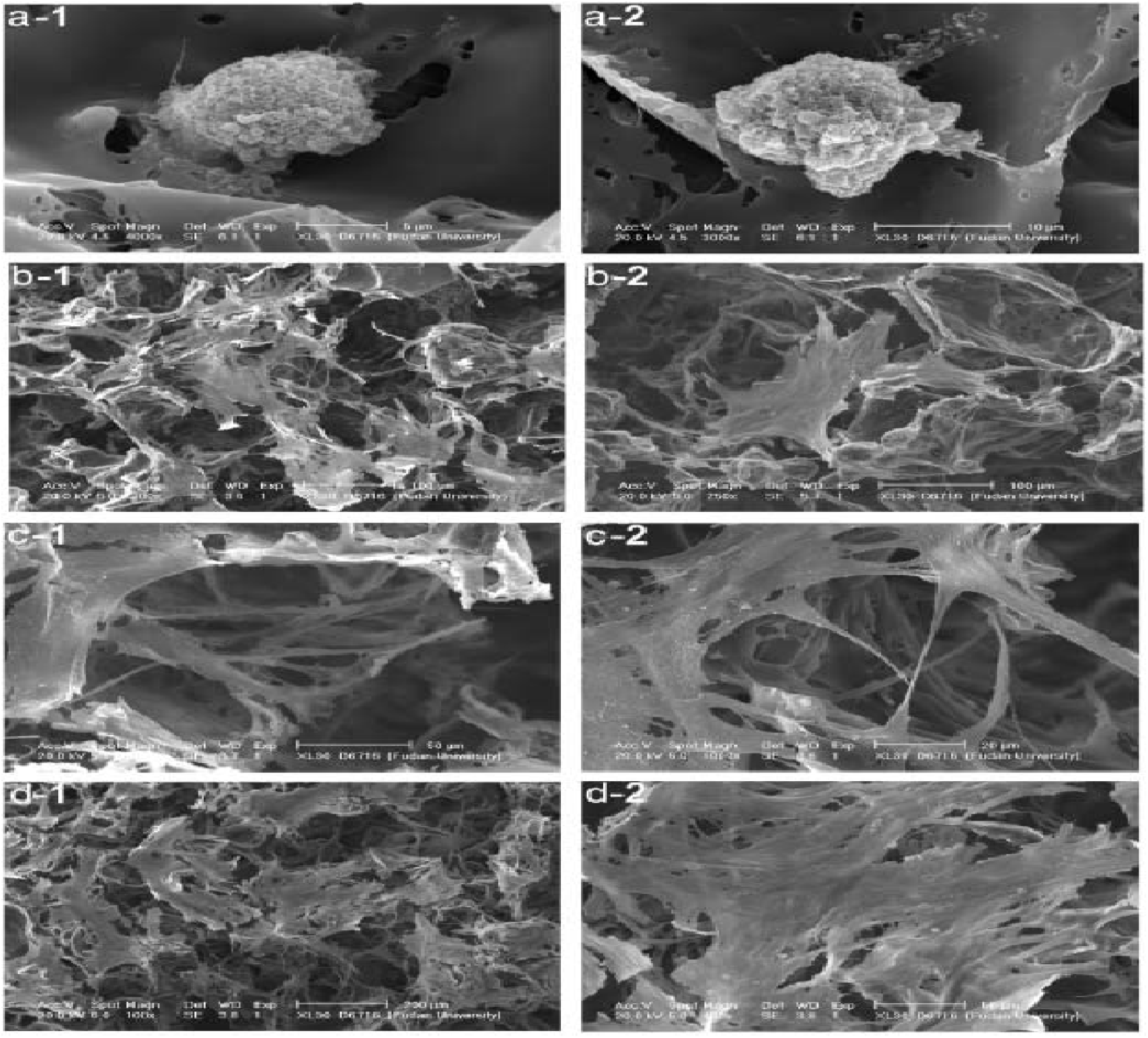
Morphological changes of fibroblast cells under SEM cultured in collagen gels (a1-d1) and changes were reordered in a2-d2 with average of cell area. The fibroblast cells with crystal violet on dish at 3 hrs.

#### Morphological and cytoplasmic domain (LCD) changes in cells of fibroblast skin of dermal

The morphological changes of fibroblast were observed in SEM with effects of cells were damaging the cells. The cell wall was distorted with loss of intercellular with incomplete cell division, and shown in the figure with A-F was seen in rough cell surface and the cytoplasmic membrane was separate from cell wall by treated the plant extracts, Additionally, the cells also showed partial to complete loss of electron density of both their cell wall and large cytoplasmic domain (LCD). Morphological changes of fibroblast were observed in SEM with cellular effects that damaged cells. The cell wall was twisted with intercellular loss by incomplete cell division and showed cell damage and the cytoplasmic membrane differed from the cell wall extracted from the plant. The cells showed a rod-shaped structure the control of antibiotic vs extracts similar results.

### The antibiotic (Chloramphenicol IV,) and compounds action on the dermal fibroblasts cells changes under scan electron microscope

The transmission electron microscopy of control (a) with uses commercial antibiotic have a fast reaction of Chloramphenicol IV,) where the figure 9 a and figure 9 b cultures of fibroblasts. Skin fibroblasts, found in control subjects or the cells were exhibit a flat morphology with multiple cytoplasmic tapering processes were seen. The oval-shaped nucleus was located in the centre of the cell, containing heterochromatin clusters near the nuclear envelope was seen in cells. The cytoplasm contains many vesicles with variable electron density, prominent Golgi structures, and mitochondria. Rister endoplasmic reticulum (RER) cisternae in fibroblasts cells appeared to be much larger in control. Some large multilamellar bodies (MLB) was commonly found in the cytoplasm of fibroblast cells. (G) Golgi structure, (M) mitochondrion, and (V) skin. Bar = 1 μm.

**Fig 8.**
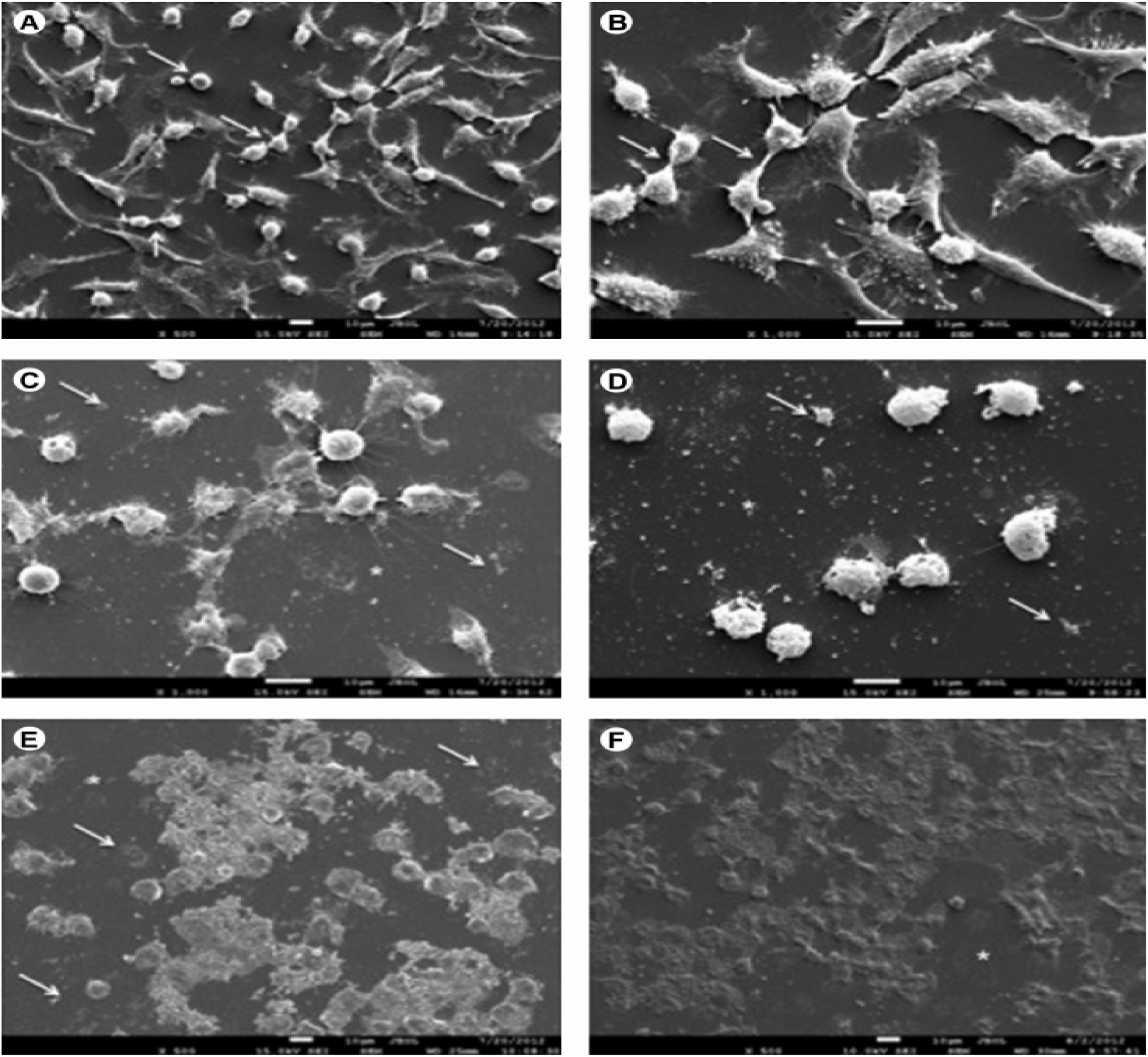
The induced morphological changes of compounds in A-F with average of cell area. The fibroblast cells with crystal violet on dish at 3 hrs. at changes

**Fig 9.**
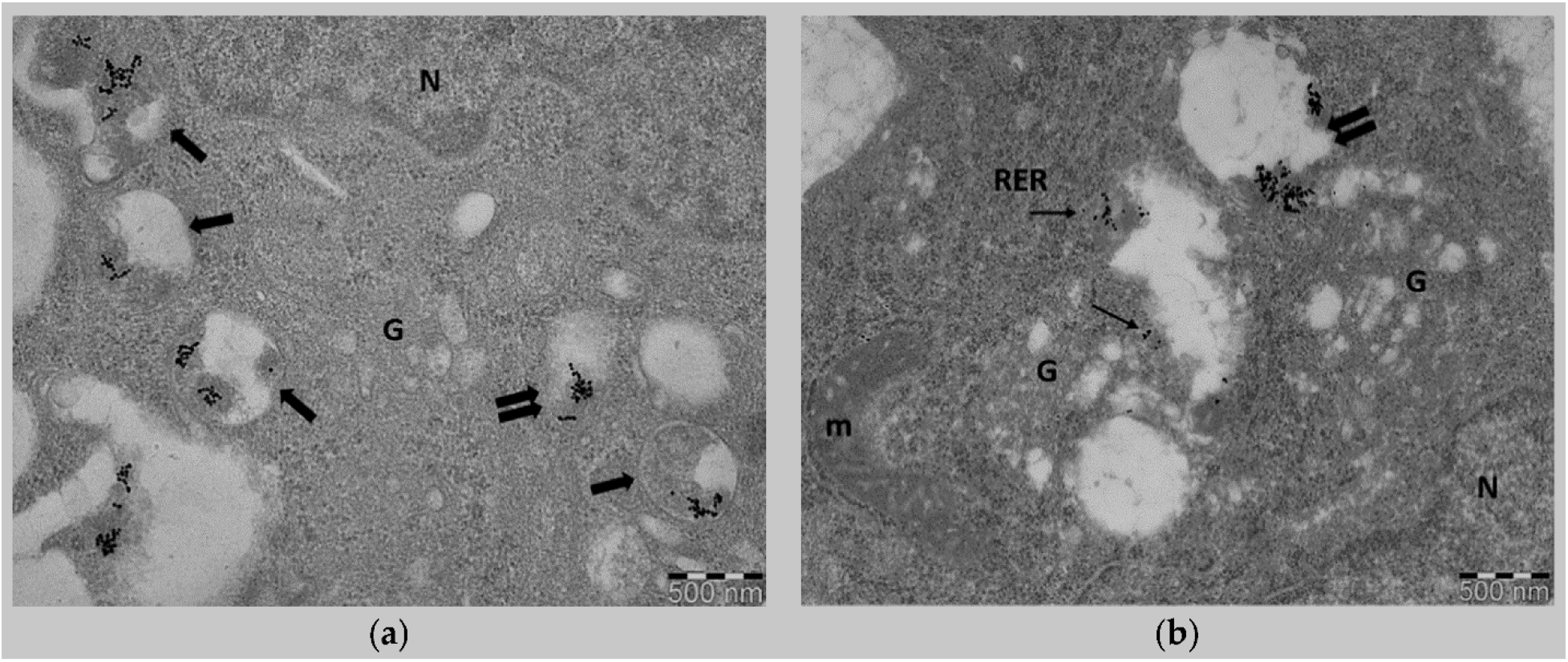
The induced morphological changes of fibroblast cells (a-b)

## 5. DISCUSSION

Continuous exposure to UV triggers the induction of images by deep tissue transformation the Collagen reduction is one of the characteristic features seen in body tissues attached to the skin created by the images. Since fibroblasts are a major source of ECM, active mutations in fibroblasts on exposed skin are considered to be related to collagen reduction. Fibroblasts often appear flat or extended on the skin and change shape to match the external environment. Previous studies have reported that most fibroblasts have a folded / heavy structure in highly damaged skin, since cell formation is a critical condition of cellular activity, the cell of fibroblast changes their morphological changes in skin fibroblasts seen in skin photography are considered related to collagen reduction there in internal cells. The advancement of the world results in the empowerment of skin disorders due to which continuous researches are in progress to control the infectious diseases with new and active regents of skin (Carvalho et al., 2005; Gulluce et al., 2006). The creams and locations with the various class of chemistry are present for the treatment of skin diseases (Ahmed et al., 2018; Khan et al., 2012). Medicinal chemists are always in search of new bioactive compounds and skin disorder agents for controlling different diseases in humans (Momtaz et al., 2008). Herbal plants have a range of potential health-related compounds (Kubo et al., 2000; Li et al., 2008). The results of current research are following the previous information where it reported that the bioactive compounds have the potential for resistance against the infectious and skin diseases of the human body (Gulluce et al., 2006). Several bioactive compounds showed antimicrobial activities at the cytoplasmic membrane by changing its function and structure and regulate infection on critical sites of cells and bind the reactive oxygen species (Natella et al., 1999). Numerous clinical researches were conducted to evaluate the different bioactive compounds for controlling numerous human diseases (Ahmed et al., 2018). The isolation of new bioactive compounds groups in the present investigation showed their mode of action for controlling the infection of diseases of skin disorder (Marra et al., 2012; Meda et al., 2005). Immunomodulation using medicinal plants can provide an alternative to conventional chemotherapy for a variety of skin diseases in the human body (Li et al., 2008). The compounds isolated from these species reflect that the cultivation of these species as a medicinally important plant will be a milestone in the medicinal world for a human being. The active compounds can act as on a sensitive nucleophilic group near the active site and the mechanism that tyrosinase regards the catechol substrate as cresol at the active site for controlling the skin diseases.

## 6. CONCLUSION

The fibroblast is a serious skin disorder the comparison of antibiotics with extracts on morphological changes the extracted cells of fibroblast showed a completed damages the cell wall and cell discretion process, the Vizic acid, psacbumin C, 8’z-enzyme conjuger, 5- (8’z, 11’z-heptodecadinyl) -1,3-benzinol, 9 ‘- (o-methyl) protocoteric acid and caliphenic acid are extracted from herbal plants as a source of novel drugs compounds were used as an antibiotic, the compounds can treat fibroblast cells of skin all compounds were chek their collagenase inhibition assay, elastase inhibition assay, hyaluronidase inhibition assay method, tyrosinase inhibition assay approach, herbal criteria collected in identification plants of completed by FRIR analysis And the results indicated that the presence of all compounds was an active source of drugs used by both species in an effective skin disease control process. The compounds were proved that have perfect novel antibiotics and are further used in creams, locations and tablets for future skin diseases control.

## Acknowledgement

The authors highly appreciate Department of Horticulture, University of Haripur and HEC, for providing Funds to complete this project.

## Authors’ contributions

WA contributed in collecting plant sample, identification and herbarium confection. Conceived and designer the experiments: WA, RA. Performed the experiments RA, AQ, AM. Analyzed the data: WA, AQ, MA Wrote the paper: WA, RA. All the authors have read the final manuscript and approved the submission.

